# Seasonal dynamics and polyphenism of butterfly communities in the coastal plains of central Western Ghats, India

**DOI:** 10.1101/2022.02.21.478808

**Authors:** Deepak Naik, R. Shyama Prasad Rao, Krushnamegh Kunte, Mohammed S. Mustak

## Abstract

Long-term socioeconomic progress requires a healthy environment/ecosystem, but anthropogenic activities cause environmental degradation and biodiversity loss. Constant ecological monitoring is therefore necessary to assess the state of biodiversity and ecological health. However, baseline data is lacking even for ecologically sensitive regions such as the Western Ghats. We looked at the seasonality and polyphenism of butterflies of the central Western Ghats to get baseline population patterns on these charismatic taxa. We recorded 43118 individuals (175 species) using fortnightly time-constrained counts for two consecutive years, and found the peak abundance (49% of the total individuals) in post-monsoon period (Oct to Jan). The seasonal abundance was correlated with the overall increase in species richness. Habitat differences were stronger than seasonality as samples clustered based on sites. Several species also displayed polyphenism with distinct distributions of wet and dry season forms. Seasonal equitability and indicator species analysis showed distinct inter-species differences in seasonality patterns. This work provides key baseline data on the seasonal dynamics of butterflies of the Western Ghats in the context of climate change and conservation, and will help in future monitoring of this ecologically sensitive region using butterflies.

## Introduction

Ecosystems and habitats are always changing, and some of these changes are fuelled by anthropogenic pressures. Therefore, periodic ecological monitoring is necessary to access the state of ecosystem and biodiversity (Lindenmayer and Likens, 2018). This requires monitoring various environmental changes and ecological taxa both at global and local levels. However, often even baseline data is lacking due to various reasons including bureaucracy, funding, and the lack of public appeal and involvement (Lindenmayer and Likens, 2018).

Butterflies as a community has a great public appeal. Further, butterflies are easy to study due to their relatively large size and conspicuousness, and well-known taxonomy (Brown, 1991). Thus, they are usually the key taxa in biodiversity monitoring and are considered as the umbrella species in nature conservation (New, 1997). Further, while butterflies perform many ecological roles, they are very sensitive to environmental changes. They react swiftly to environmental transformations such as land use dynamics and/or vegetation that affect their abundance and/or species composition (Guedes et al., 2000; Naik *et al*., 2021; Pateman *et al*., 2012; Rákosy and Schmitt, 2011). As butterflies also reflect seasonal and other natural changes, they are highly suitable for monitoring the diversity and health of ecosystems (Bonebrake *et al*., 2010; Kunte, 1997; Rao and Girish, 2007). Thus, regular assessment of butterfly populations can reveal the extent of environmental dynamics (Beaumont and Hughes, 2002; Hayes *et al*., 2009; Sreekumar and Balakrishnan, 2001).

Exploring the local abundance, community composition, and seasonality of butterflies are also important for two reasons. First, to understand the ecology of butterfly communities of a particular region; and second, to get the baseline data on this umbrella taxon from an ecologically sensitive region in the context of climate change, habitat degradation/loss, and conservation. Such studies on community dynamics are useful in effective monitoring of ecological changes caused by anthropogenic activities such as deforestation, agricultural expansion, urbanization, etc. (Gadgil, 1996; Giriraj *et al*., 2008; Purvis and Hector, 2000). In addition, climate change and environmental change have both cause and effect on each other, and the former has a disproportionately greater effect on ecologically sensitive regions (Niyogi *et al*., 2010). Butterflies are sensitive and react rapidly to climate and habitat changes (Kunte, 1997; Molina-Martínez *et al*., 2016; Padhye *et al*., 2006; Schmitt *et al*., 2021; Warren *et al*., 2001). For example, global warming and anthropogenic land use have been shown to affect butterfly species range shifts - uphill or poleward (Konvicka *et al*., 2003; Molina-Martínez *et al*., 2016; Parmesan *et al*., 1999; Pateman *et al*., 2012). Thus understanding the seasonal dynamics of butterflies as a community has a great relevance.

The Western Ghats in India is one such ecologically diverse and sensitive mountain region; but is fast undergoing socioeconomic and industrial developments, and facing environmental degradation (Jha *et al*., 2000; Jitendra, 2019; Rao and Girish, 2007). Despite being a biodiversity hotspot, systematic ecological assessment and monitoring are scares; and limited baseline information is known about the community dynamics of various taxa, including butterflies, in this region (Kunte, 1997; Kunte *et al*., 1999; Naik *et al*., 2022). Such data are useful for both ecological monitoring and prioritizing the conservation needs of the region (Gadgil, 1996).

Butterfly community dynamics, especially in relation to season and climate change, have been keenly explored (Hamer *et al*., 2005; Molleman *et al*., 2006; Schmitt *et al*., 2021). For example, seasonality and phenology of Mexican butterflies were studied using a large sample of 60,662 individuals (Pozo *et al*., 2008). Grøtan *et al*. (2012) studied 10 year seasonal cycles of 137 fruit-feeding butterfly species (20,996 individuals) in Amazonian Ecuador. British butterflies were well studied in terms of seasonality and climate (Roy *et al*., 2001). There are many studies on the seasonality of butterflies of the Indian subcontinent (Kunte *et al*., 2019; Padhye *et al*., 2006; Sengupta *et al*., 2014). For example, Tiple and Khurad (2009) explored the seasonal distribution of butterfly species (but not their abundance) in Nagpur city of central India. Likewise, Singh *et al*. (2015) studied the seasonality of butterflies of northeast India. Recently, Sharma and Sharma (2021) studied the short-term (from April to December 2018) seasonal patterns of 118 butterfly species (6384 individuals) of northern India. There are also some studies on the seasonality of butterflies from the Western Ghats region (Arun, 2003; Kunte, 1997; Kunte *et al*., 2019; Padhye *et al*., 2006; Paul and Sultana, 2020; Revathy and Mathew, 2013). However, more comprehensive studies are required, with long-term and more widespread datasets on butterfly population dynamics and seasonality. Further, seasonal dynamics of butterflies have not been explored in the coastal regions and foot hills of the Western Ghats (Naik *et al*., 2022).

Another important ecological phenomenon in many species of butterflies is the seasonal polyphenism in which distinct phenotypes occur from the same genotype (Brakefield and Larsen, 1984; Shapiro, 1976). Changing seasons or habitat conditions lead to changes in wing coloration of butterflies in successive generations, which helps them blend in with the environment and avoid predation (Brakefield, 1987; Mohandas and Remadevi, 2019). Seasonal polyphenism was investigated in Holarctic, and African nymphalids and pierids (Brakefield, 1987; Brakefield *et al*., 2007; Kato and Handa, 1992). But studies are limited in Indian subcontinent, with very few species (such as *Chilades pandava* and *Melanitis leda*) being studied in relation to population dynamics (Tiple *et al*., 2009). Therefore, exploring the seasonal polyphenism in butterflies of the Western Ghats is of great interest and relevance.

The aim of this paper was to obtain baseline data on seasonal patterns of butterfly communities of the Western Ghats - an anthropogenically dynamic and ecologically sensitive region. We used fortnightly time-constrained counts to get a large sample of 43,118 individuals from eight heterogeneous landscapes of the central Western Ghats and present the seasonal trends in diversity and species-specific abundance. We also discuss the relevance of this study in the context of ecological monitoring, climate change, and conservation.

## Materials and Methods

### Study sites

This study was conducted in eight sites/landscapes/habitats in the coastal plains and foothills (in the western side) of the central Western Ghats in the Dakshina Kannada district of Karnataka, India. Detailed descriptions of the study sites [namely Agriculture (AR), Botanical arboretum (BA), Coastal (CO), Laterite mixed shrubby (LMS), Modified forest (MF), Mixed moist deciduous (MMD), Rocky crop (RC), and Semi evergreen (SE)] and the location map were given in our previous paper (Naik *et al*., 2022). While there was no demarcation for the study sites from their surroundings, their rough area varied from 1.6 sq km (Coastal) to several hundred sq km continuum of the Western Ghats (Semi evergreen). The study sites are in elevation ranging from 4 m to 304 m. On average, the study area (Dakshina Kannada district, between 12°28’N to 13°10’N and 74°48’ to 75°40’E) receives an annual mean rainfall of 3916 mm (India Meteorological Department [IMD], https://mausam.imd.gov.in/), and has a tropical temperature/climate (see supplemental information - materials and methods). Monthly rainfall data for the area were obtained from IMD (https://mausam.imd.gov.in/), and day temperature and relative humidity data from each site were collected (using a HTC AVM-06 digital anemometer) during fortnightly field visits. No other environmental parameters have been collected. Permission for the field survey was taken from the Principal Chief Conservator of Forests, Karnataka Forest Department, Bengaluru (No. PCCF/C/GL-01/2016-17). No butterflies have been killed or collected in this study.

### Butterfly survey

The butterfly surveys were performed systematically for two consecutive years between November 2016 and October 2018 to collect the quantitative data on the abundance of butterfly communities at each site. Detailed description of the butterfly survey was presented in our previous paper (Naik *et al*., 2022). In brief, data were collected using time-constrained count method (Kadlec *et al*., 2012; Pollard, 1977; Suman *et al*., 2021) by walking a transect trail (0.5 km) slowly in 30 min during the peak activity of butterflies between 9:30 AM and 1:00 PM. Every site had six spatial replicates (consecutive transects of 0.5 km each) and was covered in a total of three hours (30 min per transect). Each site was visited systematically once in 15 days, and the butterflies present within a width of 5 meters on either side of the transect were identified on the move and noted to species level (Table S1) (Kehimkar, 2008; Kunte, 2000; Naik *et al*., 2022). However, in some cases where there was difficulty in identifying individuals to the species level in the field, they were noted only to the genus level.

We also recorded the number of individuals in wet season form (WSF) and dry season form (DSF) for five species (namely *Jamides celeno, Melanitis leda, Mycalesis* spp. [*M. mineus* or *M. perseus*, mostly the first one], *Ypthima baldus*, and *Ypthima huebneri*) with polyphenism wherein WSF and DSF could easily be recognized in the field (see supplemental information and Table S1) (Bhakare and Ogale, 2018; Kunte *et al*., 2021; Nitin *et al*., 2018).

Further, common plants present along the transect at each site were identified to the species level (a few to the genus level only) and their habits were noted (Naik *et al*., 2021; 2022). It was just an indicative presence-absence data and not a systematic or quantitative survey (Naik *et al*., 2022). The species that are potential hosts for the butterflies of the Western Ghats were noted based on the literature (Kunte et al., 2021; Naik *et al*., 2021; 2022; Nitin et al., 2018).

### Data/statistical analyses

A total of 384 days of census was carried out. After pooling fortnightly samples, there were 144 samples (6 replicates * 24 months) for each site. Due to short-term nature of the study (for only two years) and low number of individuals for many species/site-wise replicates, interannual differences were not explored and only limited site-specific analyses were performed. Thus, samples were further combined, where required/appropriate, to get monthly/seasonal patterns site-wise or overall (Table 1 and S1). While seasonal variations in diversity and abundance have been presented as mean ± standard deviation based on replicates (Table 1) for individual sites or for all sites combined; where requisite, further analyses and presentations were done using all replicates combined.

**Table 1.** Seasonal abundance of butterfly communities of the Western Ghats.

| Habitat | Species<sup>#</sup> |  |  |  | Individuals<sup>#</sup> |  |  |  |
| --- | --- | --- | --- | --- | --- | --- | --- | --- |
|  | Pre-monsoon | Monsoon | Post-monsoon | Total | Pre-monsoon | Monsoon | Post-monsoon | Total |
| **Agriculture** | 26.7±6.0<sup>a+</sup> | 34.8±4.6<sup>a</sup> | 47.7±4.8<sup>b</sup> | 55.8±7.3 | 211±107<sup>a</sup> | 203±87.5<sup>a</sup> | 502±212<sup>b</sup> | 917±383 |
| **Botanical arboretum** | 37.2±10.5<sup>a</sup> | 54.0±10.4<sup>b</sup> | 65.3±6.2<sup>b</sup> | 78.8±10.2 | 341±116<sup>a</sup> | 347±103<sup>a</sup> | 536±152<sup>a</sup> | 1224±312 |
| **Coastal** | 18.0±4.1<sup>a</sup> | 28.2±4.6<sup>b</sup> | 31.3±5.2<sup>b</sup> | 40.7±5.1 | 92.0±18.8<sup>a</sup> | 221±57.5<sup>b</sup> | 291±48.3<sup>b</sup> | 604±110 |
| **Laterite mixed shrubby** | 23.0±4.1<sup>a</sup> | 53.5±8.9<sup>b</sup> | 50.8±3.7<sup>b</sup> | 70.7±6.5 | 169±70.2<sup>a</sup> | 319±166<sup>ab</sup> | 328±67.6<sup>b</sup> | 816±273 |
| **Modified forest** | 42.2±9.8<sup>a</sup> | 50.5±8.3<sup>ab</sup> | 63.0±12.6<sup>b</sup> | 78.8±13.2 | 227±83.8<sup>a</sup> | 277±68.1<sup>a</sup> | 502±153<sup>b</sup> | 1005±262 |
| **Mixed moist deciduous** | 36.2±10.0<sup>a</sup> | 40.2±10.5<sup>a</sup> | 54.3±11.6<sup>a</sup> | 69.8±14.3 | 169±61.5<sup>a</sup> | 166±38.7<sup>a</sup> | 299±74.6<sup>b</sup> | 634±136 |
| **Rocky crop** | 23.3±6.4<sup>a</sup> | 31.5±7.1<sup>a</sup> | 51.5±6.5<sup>b</sup> | 64.5±8.9 | 165±34.2<sup>a</sup> | 129±24.8<sup>a</sup> | 291±68.6<sup>b</sup> | 584±88.6 |
| **Semi evergreen** | 52.5±8.5<sup>a</sup> | 49.3±6.6<sup>a</sup> | 69.2±4.5<sup>b</sup> | 88.0±7.2 | 422±101<sup>b</sup> | 212±73.0<sup>a</sup> | 769±214<sup>c</sup> | 1403±374 |
| **Total** | **91.7±11.4<sup>a</sup>** | **109±8.0<sup>b</sup>** | **124±3.3<sup>c</sup>** | **141±5.4** | **1795±300<sup>a</sup>** | **1874±309<sup>a</sup>** | **3517±486<sup>b</sup>** | **7186±1075** |

All values are given as mean ± SD (n = 6).
<sup>#</sup>Scheirer–Ray–Hare test results: [H = 56.3 and p = 8E-10 for habitat, H = 44.9 and p = 2E-10 for season, and H = 11.9 and p = 0.61 for habitat:season] for species and [H = 38.6 and p = 2E-6 for habitat, H = 45.3 and p = 1E-10 for season, H = 17.6 and p = 0.22 for habitat:season] for individuals.
<sup>\*</sup>Within a row and group (species or individuals), values bearing the same letter are not significantly different between seasons at p < 0.05 as determined by Kruskal-Wallis post hoc test.

Relative abundances were calculated using 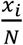 and where appropriate, normalized abundances were calculated using 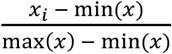(where *x* is the absolute frequency and N is the total), so that they are comparable across samples and/or species (Naik *et al*., 2022). In addition, various diversity indices, where appropriate, were calculated as explained in detail in our previous paper (Naik *et al*., 2022). In particular, seasonal equitability or evenness of species across seasons/months was calculated as Simpson’s, *E* = *D*/*D*_*max*_ where 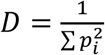, *p*_*i*_ is the proportion of *i*^th^ species, and *D*_*max*_ is the number of seasons/months (Magurran, 1988; Naik *et al*., 2022), and plotted against normalized abundance.

Seasonal patterns were visualized using trend-lines, radar charts, and/or heat maps and hierarchical clustering analysis (HCA). Relationship between seasonal species richness and abundance, and among species were explored using correlation. The HCA used Bray-Curtis dissimilarity (a measure of β-diversity) as the distance and Ward’s method as the clustering algorithm. The heat map and HCA were done using heatpmap.2() in gplots package in R software (https://cran.r-project.org/web/packages/gplots/index.html).

A non-metric multidimensional scaling (NMDS) was used to explore the patterns of similarities among months (seasons), sites (habitats), and/or species (Naik *et al*., 2022; Pozo *et al*., 2008). NMDS is a rank-based/non-parametric approach and better (than PCA, for example) when the data matrix contains a lot of zero values (Legendre and Gallagher, 2001). It was performed using Bray-Curtis dissimilarity (a measure of β-diversity) - a semimetric/abundance-based index (Greenacre and Primicerio, 2013; Schroeder and Jenkins, 2018). NMDS was done in R version 3.6.2. The number of reduced dimensions in NMDS was k = 3 such that NDMS stress was kept well below 0.15 (Naik *et al*., 2022). Visualization was done by plotting the NMDS scores in Microsoft Excel.

Finally, we also looked at the indicator values (Barrow and Parr, 2008; De Cáceres *et al*., 2010; Naik *et al*., 2022; Sharma *et al*., 2020) of butterfly species in relation to seasons. The indicator values (IndVal) were based on all possible combinations of groups of seasons (or samples/transects) and selecting the combination for which the species can be best used as an indicator (De Cáceres *et al*., 2010). The indicator value estimation was done using multipatt() in R indicspecies package which also gives p-values based on permutation test (De Cáceres *et al*., 2010). A default number of 999 permutations was used (Naik *et al*., 2022).

Where appropriate, distribution-free non-parametric statistical analyses/tests were performed (Naik *et al*., 2022). For example, Spearman’s rank correlation, which is less sensitive to outliers, was used to see the relationship among species based on seasonality. The significance of correlation coefficient was testing using cor.test() [which is based on t-distribution or approximation] in R. A chi-squared test for trend in proportions [prop.trend.test() in R, https://rdrr.io/cran/rstatix/man/prop_trend_test.html] was performed to see if two groups followed a specified trend (for example, wet and dry season forms in a particular seasonal order). A Scheirer–Ray–Hare test was performed using scheirerRayHare(y ∼ habitat + season) in R for testing the significance of observed differences in the individual and species richness among sites and seasons based on spatial replication data (Barrow and Parr, 2008; Sokal and Rohlf, 1995). Further, a Kruskal-Wallis post hoc test was performed using pairwise.wilcox.test() [with p.adjust.method=“BH”]. All statistical analyses, where explicitly not mentioned, were performed in R version 3.6.2.

## Results

### Seasonality of species richness and abundance

A total of 43,118 individuals (175 species) were recorded in this study. Detailed results on site-specific species richness and abundance were presented in a previous paper elsewhere (Naik *et al*., 2022). Seasonal pattern of species richness and abundance is presented here in Table 1 and Fig. 1A. Species richness varied from 71 to 122 per month (40.6% to 69.7% of the maximum 175 species, 53.7% to 77.1% species if two years combined) - highest in early post-monsoon (Oct to Jan).. The overall (two years combined) monthly abundance pattern is shown as a radar plot (Fig. 1B) wherein 49% of individuals occur in post-monsoon period (Fig. 1C). Seasonal trend of individual richness was correlated to species richness (ρ = 0.771, p = 1.0E-5, t-test for correlation, Fig. 1A and D) and it was more pronounced (ρ = 0.86, p = 1.4E-7) if an outlier (May 2007 data point) was ignored. The monthly patterns of butterflies of the Western Ghats is presented in Table S1. Table S2 presents taxonomic family-wise distribution of species richness and abundance of butterflies in three seasons. Being the most abundant in terms of individuals, Nymphalidae had a seasonal trend more similar to the total (χ2 = 6.0, p = 0.014, χ^2^ test for trend in proportions), but other families had very dissimilar seasonal trends. Abundances of Hesperiidae, Lycaenidae, and Nymphalidae increased by 663, 183, and 99%, respectively, from pre-monsoon to post-monsoon, whereas Papilionidae and Pieridae abundances were more similar in three seasons. A summary of some key diversity indices is shown in Table S3. Overall, dry season has lower diversity which was very pronounced (for example, in terms of rarefied richness and Shannon’s and Simpson’s indices) in May.

**Fig. 1.**
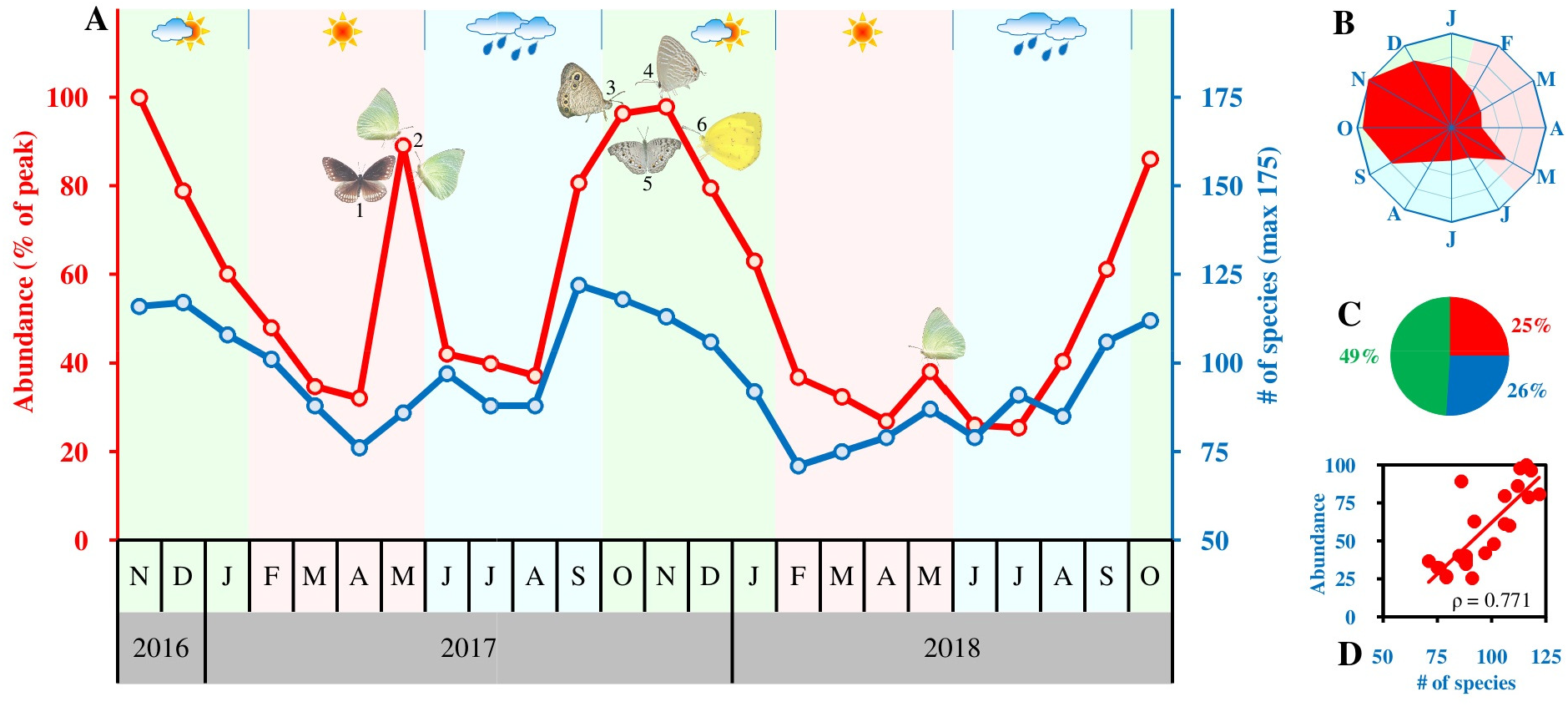
Seasonality of butterflies of the Western Ghats. (A) Plot of abundance (% of peak in Nov 2016 with 3189 individuals) and the species richness of butterflies. Butterfly abundance peaks in post-monsoon period (Oct to Jan) and lowest abundance was at 25.3% of the peak. Monthly number of species varied from 71 to 122 (40.6% to 69.7% of the maximum). May 2017 showed a peak due to odd migration event of *Catopsilia* spp. (B) Radar plot of pooled abundance mainly occupies upper-left part - in the post-monsoon period. (C) The pie chart shows percentage of abundance in three periods with post-monsoon (green sector) having nearly 49% of the total annual number of butterflies. (D) Strong positive correlation (p = 1.0E-5) is seen between abundance and species richness. Flagged (some of the abundant) species in the figure are (1) *Euploea* spp., (2) *Catopsilia* spp., (3) *Ypthima huebneri*, (4) *Jamides celeno*, (5) *Junonia atlites*, and (6) *Eurema hecabe*.

There was difference in the overall pattern of both species richness and abundance between two years of sampling (Fig. 1A). One obvious difference, but can be considered as an anomaly, was a very pronounced peak in the abundance of *Catopsilia* spp. (mainly *C. pomona*) in May 2017 compared to May 2018 (see next section and supplemental information - results and discussion).

### Seasonality versus site-specificity

NMDS plot (Fig. 2A) clearly showed samples in a circular order according to the month of sampling. There were differences between two years of sampling, and May 2017 was the obvious outlier. However, plot did not reveal distinct groupings of samples according to three seasons (pre-monsoon, monsoon, and post-monsoon). When site-wise samples were considered, NMDS plot (Fig. 2B) showed samples clustering based on sites rather than month of sampling. For example, monthly samples from coastal (CO) habitat formed a clear cluster. However, when there were overlaps among sites, samples from the same months/season (for example, May) appeared to occur nearby.

**Fig. 2.**
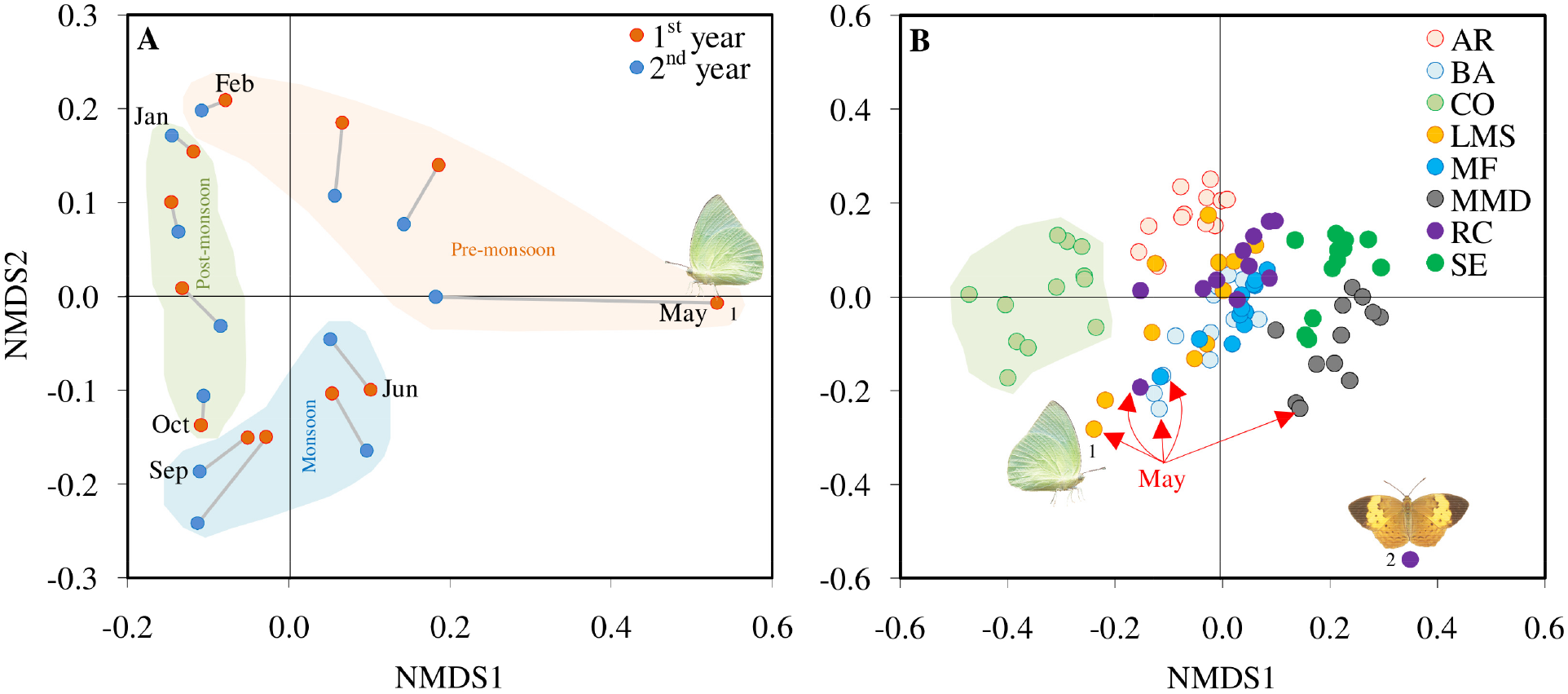
Seasonality and site-specificity using NMDS. (A) The NMDS plot (stress = 0.05) shows samples in a circular order according to the month of sampling. May sample was an outlier due to overabundance/migration event of *Catopsilia* spp. in the year 2017. (B) Clustering is stronger based on sites compared to month of sampling; however, when sites overlap, samples from same month seem to occur nearby (NMDS stress = 0.135). An obvious outlier (rocky crop [RC], July) in the plot was due to very small sample size (N = 4) in which 50% were *Cupha erymanthis*. Flagged species in the figure are (1) *Catopsilia* spp. and (2) *Cupha erymanthis*.

Although overall species richness and abundance peaked in post-monsoon period (Fig. 1A-C), and site-specific patterns consistently reflected that general trend (Table 1); there were large differences among sites/habitats. Both habitat and season influenced species and individual richness. However, seasonal effect was much stronger for individual richness (H = 38.6 and p = 2E-6 for habitat versus H = 45.3 and p = 1E-10 for season, Scheirer–Ray–Hare test). The habitat:season interaction was not significant. A HCA and heat map of site-wise relative monthly abundances of 54 species of butterflies (that together make up more than 90% of the total) is shown in Fig. S2. It is clear that site-specific differences in the abundance were more pronounced than seasonality as samples mainly clustered based on sites (habitats) rather than months of sampling. For example, most of the samples from agriculture (AR) or mixed moist deciduous (MMD) habitats cluster together. This is because different habitats showed large differences in the composition of species (Naik *et al*., 2022).

### Interspecies variation in the seasonality

Fig. 3A presents HAC and heat map of overall relative monthly abundances of 54 species of butterflies (that together make up more than 90% of the total abundance). While species diversity and abundance were highest at late monsoon and early post-monsoon season (Fig 1A), there was great seasonal variation in the abundance of different species (Fig. 3A and B). For example, the most abundant *Euploea* spp. (n = 3080, mainly *E. core*) occurred mostly in dry season (pre-monsoon, Feb to May) peaking at Apr. Further, it showed similar pattern of occurrence in two years of sampling as against *Catopsilia* spp., the second most-abundant (n = 2752) that peaked in May (Fig. 3B). While many species, such as *Leptosia nina* (n = 557) were found more uniformly throughout the year, species such as *Prosotas dubiosa* (n = 324) showed very clear bimodal seasonal abundance. Relationship between the relative abundance and seasonal equitability (a measure of evenness) of butterflies is shown in Fig. 3C. It is evident that seasonal evenness increases as abundance increases (ρ = 0.807, p = 1.9E-41). However, many species are seasonally more even in spite of their low abundance or vice versa (for example, *Catopsilia* spp.). Fig. S3 shows the heat map and HCA of Spearman’s rank correlation of butterflies wherein species clustered based on seasonality. Interspecies correlation coefficient ranged from -0.928 to 0.965.

**Fig. 3.**
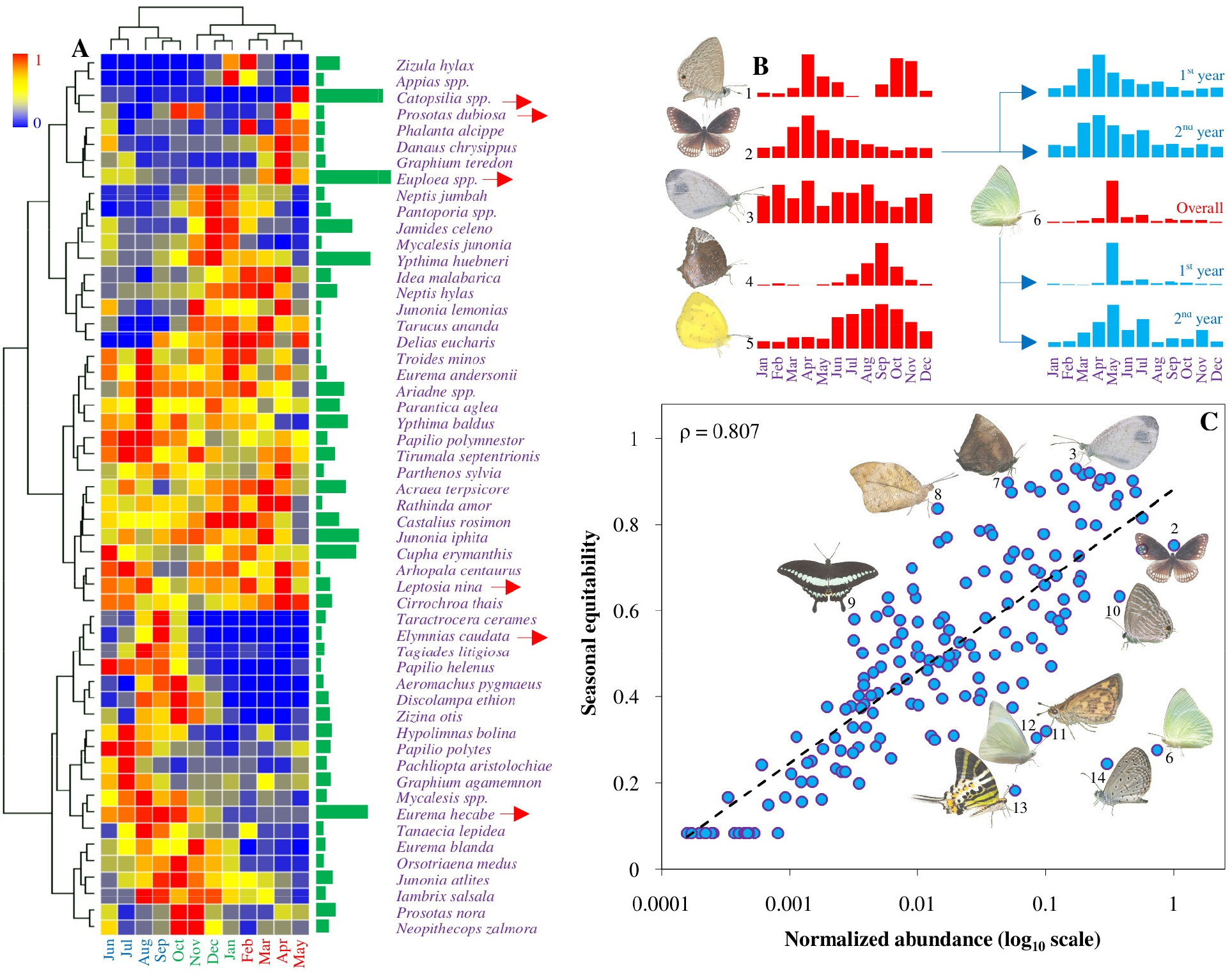
Seasonality of different species of butterflies of the Western Ghats. (A) HCA and heat map of top 54 species that make up more than 90% of the total individuals (green bars next to heat map show total abundance). Many species are abundant only in a few months. (B) Bar chart of abundance of some select species (red arrows in A) with distinct seasonality patterns. Inter-annual pattern is similar in *Euploea* spp., but very different in *Catopsilia* spp. (C) Plot of seasonal equitability against relative abundance. Evenness of butterfly species across seasons is correlated (ρ = 0.807) with overall abundance, but many species which were less abundant showed high evenness and vice versa. Flagged species in the figure are (1) *Prosotas dubiosa*, (2) *Euploea* spp., (3) *Leptosia nina*, (4) *Elymnias caudata*, (5) *Eurema hecabe*, (6) *Catopsilia* spp., (7) *Arhopala centaurus*, (8) *Hebomoia glaucippe*, (9) *Papilio liomedon*, (10) *Jamides celone*, (11) *Taractrocera ceramas*, (12) *Appias* spp., (13) *Graphium antiphates*, and (14) *Zizula hylax*.

An NMDS of species (Fig. S4) showed a pattern according to their abundance. Although many abundant species had distinct seasonality patterns (Fig. 3 and S2), they clustered closely together at the center of NMDS plot. Less abundant species were more spread apart, and the singletons were at the periphery of the plot. Further, singletons showed a circular order according to the month of occurrence. It is obvious that singletons exerted a distorting effect on the NMDS plot as the single individual can occur only in one sample/month. However, abundant species with strong seasonality (for example: *Catopsilia* spp. in May, *Discolampa ethion* in Oct-Nov, and *Zizula hylax* in Jan-Feb) nearly followed the circular order of singletons.

### Seasonal indicator species

There were 38 seasonal indicator species (or 78 species when taken on monthly basis) (Table S4). While monsoon and post-monsoon seasons had numerous common indicator species, there were only 11 single-season-specific indicator species and a few more species in combination with pre-monsoon (dry) season (Fig. S5). For example, *Graphium antiphates* was found exclusively in pre-monsoon (dry) season (p < 0.05, permutation test), while *Kallima horsfieldii* and *Doleschallia bisaltide* mostly found in monsoon season. A few more indicator species such as *Graphium doson* and *Appias* spp. were found in two seasons in combination with pre-monsoon (dry) season. The proportion of indicator species ranged from 0.23 (April/July) to 0.49 (November) when taken month-wise, and 0.07, 0.15, and 0.23 for pre-monsoon, monsoon, and post-monsoon seasons, respectively. It may be noted that *Catopsilia* spp., while overabundant in May, was not found to be an indicator species for any season.

### Seasonal polyphenism

Patterns of seasonal polyphenism are shown in Fig. 4A-E (radar plots in the top panel) for five species (namely *Jamides celeno, Melanitis leda, Mycalesis* spp., *Ypthima baldus*, and *Ypthima huebneri*) wherein individuals could easily be identified as WSF or DSF in the field based on their distinct wing markings (Fig. 4A-E, image pairs in the middle panel - WSF and DSF, respectively). From the radar plots, it is very clear that WSF and DSF had very distinct/non-overlapping distributions. The WSF peaked between Oct to Dec (early to mid-post-monsoon period) and DSF peaked between Dec to Feb (mid post-monsoon to early pre-monsoon period) - the latter trailing, on average, by at least two months. However, in *Melanitis leda* (χ^2^ = 7.7, p = 0.0055, χ^2^ test for trend in proportions) and more strongly in *Mycalesis* spp. (χ^2^ = 0.35, p = 0.55), there was a shift in the WSF toward monsoon and DSW toward post-monsoon (Table S5). It should be noted that the proportions of DSF were much lower (12.1 to 45%) than WSF (Fig. 4A-E, pie charts in the bottom panel and Table S5). *Melanitis leda* showed a relatively high proportion of DSF (45%), but was the least abundant (N = 151) among the polyphenic species studied here (see supplemental information - results and discussion). Annual and site-specific variations in the polyphenism were not explored in-depth due to small sample sizes.

**Fig. 4.**
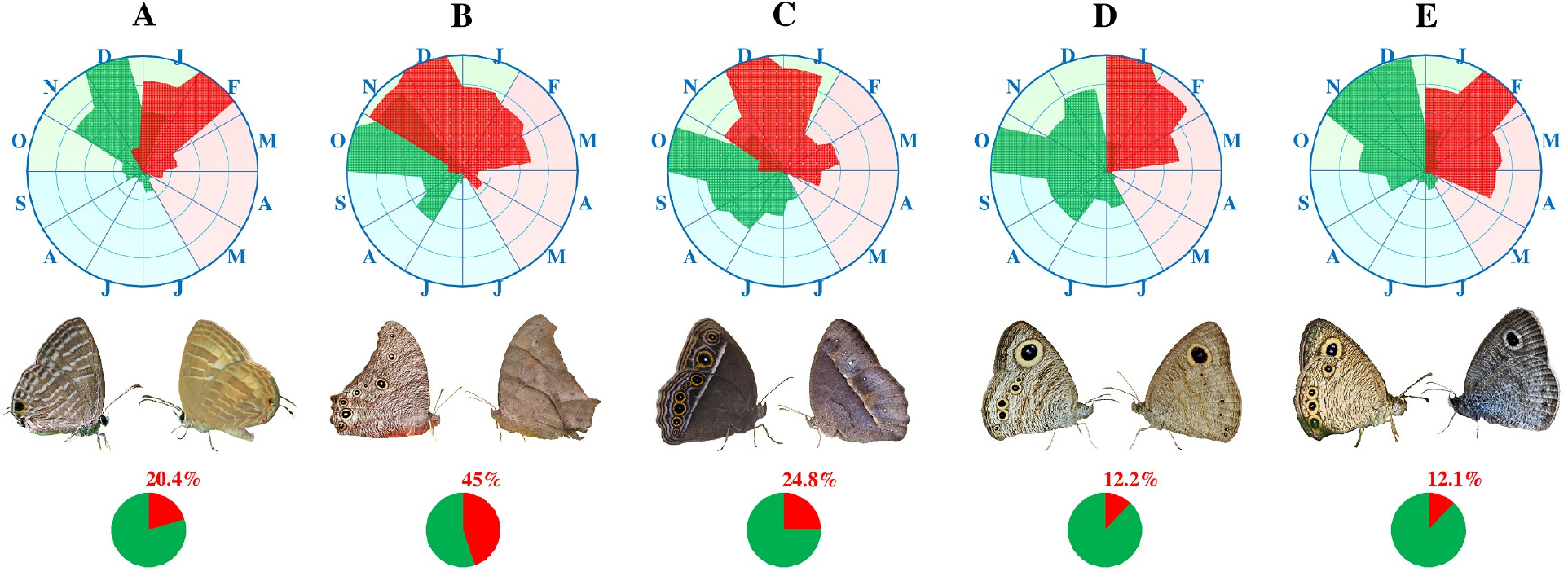
Polyphenism of butterfly species in the Western Ghats. Filled radar plots showing the abundance of wet season forms (WSF) and dry season forms (DSF) of (A) *Jamides celeno* (N = 1475), (B) *Melanitis leda* (N = 151), (C) *Mycalesis* spp. (N = 427), (D) *Ypthima baldus* (N = 1286), and (E) *Ypthima huebneri* (N = 2249). In all species WSF peaks in Oct-Dec and DSF in Dec-Feb, and the DSF trails the WSF by at least two months. Middle panel shows representative photos of WSF and DSF, respectively, for each species. Lower panel shows the percentage of DSF.

## Discussion

Many studies have explored the seasonal patterns of butterfly communities in different region of the world (Hamer *et al*., 2005; Molleman *et al*., 2006; Schmitt *et al*., 2021). There are several studies on the distribution and composition of butterflies of the Western Ghats (see a list in Padhye *et al*., 2012). Starting with Kunte (1997), there are some systematic studies on the seasonality of butterflies in India (Paul and Sultana, 2020; Sengupta *et al*., 2014; Sharma and Sharma, 2021; Singh *et al*., 2015), and in particular, in the Western Ghats (Chandekar *et al*., 2014; Mohandas and Remadevi 2019; Padhye *et al*., 2006). In this work, using a large sample, we explored the seasonal dynamics of butterfly communities in diverse habitats among the coastal plains and foot hills of the central Western Ghats.

Overall, both species diversity and abundance peaked in late monsoon and early post-monsoon season. Several factors such as day length, temperature, humidity, and precipitation greatly influence the seasonal abundance and activity of organisms both directly, and indirectly through food resources (Shimadzu *et al*., 2013; Tiple *et al*., 2009). In the Western Ghats, monsoon rains lead to lush green vegetation (especially herbaceous and grass species) in the late monsoon and post-monsoon period that provide an overabundance of host resources to the majority of butterfly species (Naik *et al*., 2021). On the other hand, seasonal leaf flushes of larval hosts such as *Cassia* and *Senna* spp., might be a key reason for the peak of *Catopsilia* spp. in May (Bhaumik and Kunte, 2020). Likewise, many species might also depend on specialist needs such as specific phenology (Navarro-Cano *et al*., 2015) or habitats (Dilts *et al*., 2019). For example, *Graphium antiphates* predominantly occurs in pre-monsoon, and *Kallima horsfieldii* in early monsoon season. Similarly, *Taractrocera ceramas* occurs mostly in monsoon season, possibly because it mainly depends on host *Oryza sativa* (Naik *et al*., 2021) which is being cultivated in monsoon season. However, as plant phenology is subject to fluctuations from numerous factors such as monsoon, microclimate, and local geography (Montgomery *et al*., 2020), seasonality of butterflies are also subject to inter-annual fluctuation as seen here, especially so for *Catopsilia* spp. and *Euploea* spp. (see supplemental information - results and discussion).

Seasonality might also depend on other factors such as migration, mimicry/inter-species interactions, and predation (Bhaumik and Kunte, 2018; Finkbeiner *et al*., 2018). For example, species in the genus *Graphium* which all have overlapping host in Annonaceae, Lauraceae, and Magnoliaceae (Naik *et al*., 2021) have quite distinct seasonal distributions - *Graphium agamemnon* dominates in monsoon and post-monsoon, *Graphium antiphates* is exclusively found in pre-monsoon, while *Graphium doson* and *Graphium teredon* dominate in pre-monsoon and monsoon seasons. It was also said that some species/populations maintain low numbers during monsoon to avoid heavy rainfall (Kunte, 1997), while some others such as *Euploea* spp. and *Tirumala* spp. start migration to plains to escape from southwest monsoon (Bhaumik and Kunte, 2018). Extent of seasonality might also depend on the species traits. For example, large sized butterflies are known to show stronger seasonality (Ribeiro and Freitas, 2011).

Another seasonal trait - polyphenism, with its ecological cause and effect - is strongly influenced by various environmental factors (van Bergen *et al*., 2017). Apart from a long list of species, very little is known about the polyphenic butterflies of the Western Ghats (Halali *et al*., 2021; Molleman *et al*., 2020; Tiple *et al*., 2009). This work provides data on the relative seasonal proportions of numerous polyphenic butterfly species of the Western Ghats, and shows distinct distribution patterns of wet and dry season forms for a select few. It may be noted that majority of the individuals were of wet season form. The changes in the proportions of wet and dry season forms and the extent of seasonal shifts as the Western Ghats experience a fluctuating or declining monsoon will be interesting topics to explore.

Finally, systematic surveys like this have a great relevance (for example, based on reliability of data and availability of baseline information) for ecological monitoring and conservation, and for studies on habitat and climate change (Cohn, 2008; Mitchell *et al*., 2017; Prudic *et al*., 2017). As the climate and Indian summer monsoon rainfall are changing over different regions (Varikoden *et al*., 2019), it would be important to record the baseline data and to track the extent of seasonal shifts in the butterfly communities of the Western Ghats. Agricultural intensification and land use change have also known to reduce or alter the monsoon rainfall (Niyogi *et al*., 2010). Climate change is also known to reduce the species diversity (Midgley *et al*., 2002) which might adversely affect butterflies directly and indirectly as they depend on a range of plant hosts (Naik *et al*., 2021). Habitat and climate changes might also influence monsoon-driven butterfly migrations (Bhaumik and Kunte, 2018; 2020).

The focus on small geographical area (central Western Ghats), short duration (two years), and non-inclusion of environmental/other factors driving seasonality are some of the limitations of this study. However, as good/systematic datasets are scarce and difficult to obtain, this work provides an extremely valuable large-scale baseline dataset on the seasonality/community dynamics which might help monitor the ecological health of butterfly populations in the Western Ghats, and could inspire other monitoring projects.

In conclusion, we looked at the seasonality and polyphenism of butterfly communities of the central Western Ghats. Using a large sample of 43,118 individuals (175 species), we showed that abundance peaked at post-monsoon period (Oct to Jan) that accounted for 49% of the total individuals. This increase was correlated with the increase in species richness. Habitat differences were stronger than seasonality as samples clustered based on sampling sites. Species showed distinct seasonality patterns as evident from seasonal equitability and indicator species analysis. Many species showed polyphenism with distinct distributions of wet and dry season forms, latter trailing by about two months. This study gives important baseline data on the seasonality of the butterfly communities of the Western Ghats, and will help in future monitoring of this ecologically sensitive region using butterflies as indicator taxa.

## Acknowledgments and Funding

Authors thank Deviprasad KN (Vivekananda College, Puttur), Dipendra Nath Basu (NCBS, Bangalore), and Nagarjuna Pasupuletti (Mangalore University) for informal discussion and suggestions on the work, Abhishek, Nishanth Katta, Vinod Simon Pinto, and Vivek Hasyagar for help in fieldwork, and Dr. Kiran Kiggal for providing some butterfly photos. Authors also thank the Karnataka Forest Department, Bengaluru, and the Pilikula Biological Park, Mangaluru for providing permissions (No. PCCF/C/GL-01/2016-17) for the field survey. DN thanks UGC-SAP, Department of Applied Zoology, Mangalore University for the facilities and Mangalore University - SC/ST cell for providing the fellowship.

## Statement of Ethics

The work is in compliance with ethical standards. No ethical clearance was necessary. No butterflies have been killed or collected in this study.

## Conflicts of Interest

The authors declare that there is no conflict of interest regarding the publication of this article.

## Data Availability

The data summaries are given in supplemental Tables S1. Full dataset may be obtained from the authors for collaborative work upon request.

## Author Contributions

DN and MSM initiated the work. DN collected the data. DN and RSPR analyzed the data and wrote the paper. All authors contributed intellectually, and were involved in the writing as well as revising the manuscript.

## Supplemental Information

Supplemental information for this article is available online.

## Notes

### Competing Interest Statement

The authors have declared no competing interest.

## References

Arun PR (2003). Butterflies of Siruvani forests of Western Ghats with notes on their seasonality. Zoos’ Print Journal 18:1003–1006.

Barrow L, Parr CL (2008). A preliminary investigation of temporal patterns in semiarid ant communities: Variation with habitat type. Austral Ecology 33:653–662.

Beaumont LJ, Hughes L (2002). Potential changes in the distributions of latitudinally restricted Australian butterfly species in response to climate change. Global Change Biology 8:954–971.

Bhakare M, Ogale H (2018). A guide to the butterflies of Western Ghats (India): Includes butterflies of Kerala, Tamilnadu, Karnataka, Goa, Maharashtra, and Gujarat states. Milind Bhakare (privately published). 496 pages.

Bhaumik V, Kunte K (2018). Female butterflies modulate investment in reproduction and flight in response to monsoon-driven migrations. Oikos 127:285–296.

Bhaumik V, Kunte K (2020). Dispersal and migration have contrasting effects on butterfly flight morphology and reproduction. Biology Letters 16:20200393.

Bonebrake TC, Ponisio LC, Boggs CL, Ehrlich PR (2010). More than just indicators: A review of tropical butterfly ecology and conservation. Biological Conservation 143:1831–1841.

Brakefield PM (1987). Tropical dry and wet season polyphenism in the butterfly Melanitis leda (Satyrinae): Phenotypic plasticity and climatic correlates. Biological Journal of the Linnean Society 31:175–191.

Brakefield PM, Larsen TB (1984). The evolutionary significance of dry and wet season forms in some tropical butterflies. Biological Journal of the Linnean Society 22:1–12.

Brakefield PM, Pijpe J, Zwaan BJ (2007). Developmental plasticity and acclimation both contribute to adaptive responses to alternating seasons of plenty and of stress in Bicyclus butterflies. Journal of Bioscience 32:465–475.

Brown Jr KS (1991). Conservation of neotropical environments: Insects as indicators. In Collins NM, Thomas JA (Eds), The Conservation of Insects and their Habitats, Academic Press, London. pp 349–404.

Burrow L, Parr CL (2008). A preliminary investigation of temporal patterns in semiarid ant communities:Variation with habitat type. Austral Ecology 33:653–662.

Chandekar SK, Nimbalkar RK, Kuvalekar AA (2014). The seasonal patterns in the abundance of butterflies, their biotopes and nectar food plants from Maval Thahsil, Pune district, Maharastra, India. International Journal of Plant, Animal and Environmental Sciences 4:50–64.

Checa MF, Levy E, Rodriguez J, Willmott K (2019). Rainfall as a significant contributing factor to butterfly seasonality along a climatic gradient in the neotropics. BioRxiv 630947v1.

Cohn JP (2008). Citizen science: Can volunteers do real research? BioScience 58:192–197.

De Cáceres M, Legendre P, Moretti M (2010). Improving indicator species analysis by combining groups of sites. Oikos 119:1674–1684.

Dilts TE, Steele MO, Engler JD, et al. (2019). Host plants and climate structure habitat associations of the western Monarch butterfly. Frontiers in Ecology and Evolution 7:188.

Finkbeiner SD, Salazar PA, Nogales S, et al. (2018). Frequency dependence shapes the adaptive landscape of imperfect Batesian mimicry. Proceedings of the Royal Society B 285:20172786.

Gadgil M (1996). Documenting diversity: An experiment. Current Science 70:36–44.

Gezon ZJ, Lindborg RJ, Savage A, Daniels JC (2018). Drifting phenologies cause reduced seasonality of butterflies in response to increasing temperatures. Insects 9:174.

Giriraj A, Irfan-Ullah M, Murthy MS, Beierkuhnlein C (2008). Modelling spatial and temporal forest cover change patterns (1973-2020): A case study from south Western Ghats (India). Sensors 8:6132–6153.

Greenacre M, Primicerio R (2013). Multivariate analysis of ecological data. Fundación BBVA. 336 pages.

Grøtan V, Lande R, Engen S, Sæther B-E, DeVries PJ (2012). Seasonal cycles of species diversity and similarity in a tropical butterfly community. Journal of Animal Ecology 81:714–723.

Guedes RNC, Zanuncio TV, Zanuncio JC, Medeiros AGB (2000). Species richness and fluctuation of defoliator Lepidoptera populations in Brazilian plantations of Eucalyptus grandis as affected by plant age and weather factors. Forest Ecology and Management 137:179–184.

Hamer KC, Hill JK, Mustaffa N, Benedick S, Sherratt TN, Chey VK, Maryati M (2005). Temporal variation in abundance and diversity of butterflies in Bornean rain forest: Opposite impacts of logging recorded in different seasons. Journal of Tropical Ecology 21:417–425.

Hayes L, Mann DJ, Monastyrskii AL, Lewis OT (2009). Rapid assessments of tropical dung beetle and butterfly assemblages: Contrasting trends along a forest disturbance gradient. Insect Conservation and Diversity 2:194–203.

Halali S, Halali D, Barlow HS, Molleman F, Kodandaramaiah U, Brakefield PM, Brattstrom O (2021). Predictability of temporal variation in climate and the evolution of seasonal polyphenism in tropical butterfly communities. Journal of Evolutionary Biology 34:1362–1375.

Holland SM (2003). Analytic Rarefaction 1.3. https://strata.uga.edu/software/ (last accessed on 13-01-2022).

Jha CS, Dutt CBS, Bawa KS (2000). Deforestation and land use changes in Western Ghats, India. Current Science 79:231–238.

Jitendra (2019). Western Ghats at risk: Deforestation data drives home point again. Down to Earth (10 May 2019). https://www.downtoearth.org.in/news/forests/western-ghats-at-risk-deforestation-data-drives-home-point-again-64470 (last accessed on 13-01-2022).

Kadlec T, Tropek R, Konvicka M (2012). Timed surveys and transect walks as comparable methods for monitoring butterflies in small plots. Journal of Insect Conservation 16:275–280.

Kato Y, Handa H (1992). Seasonal polyphenism in a subtropical population of Eurema hecabe (Lepidoptera, Pieridae). Japanese Journal of Entomology 60:305–318.

Kehimkar I (2008).The Book of Indian Butterflies. Bombay Natural History Society, Oxford University Press, Mumbai, 497 pages.

Konvicka M, Maradova M, Benes J, et al. (2003). Uphill shifts in distribution of butterflies in the Czech Republic: effects of changing climate detected on a regional scale. Global Ecology and Biogeography 12:403–410.

Kunte K (1997). Seasonal patterns in butterfly abundance and species diversity in four tropical habitats in northern Western Ghats. Journal of Bioscience 22:593–603.

Kunte K (2000). India, a lifescape: Butterflies of Peninsular India. Indian Academy of Sciences, Bangalore, and University Press, 270 pages.

Kunte K, Basu DN, Kumar GG (2019). Taxonomy, systematics, and biology of Indian butterflies in the 21st century. Indian Insects, CRC Press, London. pp 275–304.

Kunte K, Joglekar A, Utkarsh G, Padmanabhan P (1999). Patterns of butterfly, bird and tree diversity in the Western Ghats. Current Science 77:577–586.

Kunte K, Sondhi S, Roy P (2021). Butterflies of India, v. 3.11. Indian Foundation for Butterflies (https://www.ifoundbutterflies.org/).

Legendre P, Gallagher ED (2001). Ecologically meaningful transformations for ordination of species data. Oecologia 129:271–280.

Lindenmayer DB, Likens GE (2018). Effective ecological monitoring. CSIRO publishing, Australia. 210 pages.

Magurran AE (1988). Ecological diversity and its measurement. Princeton University Press, Princeton, NJ. 179 pages.

Midgley GF, Hannah L, Millar D, Rutherford MC, Powrie LW (2002). Assessing the vulnerability of species richness to anthropogenic climate change in a biodiversity hotspot. Global Ecology and Biogeography 11:445–451.

Mitchell N, Triska M, Liberatore A, Ashcroft L, Weatherill R, Longnecker N (2017). Benefits and challenges of incorporating citizen science into university education. PLoS ONE 12:e0186285.

Mohandas TV, Remadevi OK (2019). Species diversity and distribution of butterflies in Kudremukh national park and Mookambika and Someshwara wildlife sanctuaries in central Western Ghats of Karnataka. Annals of Entomology 37:113–125.

Molina-Martínez A, León-Cortés JL, Regan HM, Lewis OT, Navarrete D, Caballero U, Luis-Martínez A (2016). Changes in butterfly distributions and species assemblages on a Neotropical mountain range in response to global warming and anthropogenic land use. Diversity and Distributions 22:1085–1098.

Molleman F, Halali S, Kodandaramaiah U (2020). Brief mating behavior at dawn and dusk and long nocturnal matings in the butterfly Melanitis leda. Journal of Insect Behavior 33:138–147.

Molleman F, Kop A, Brakefield PM, DeVries PJ, Zwaan BJ (2006). Vertical and temporal patterns of biodiversity of fruit-feeding butterflies in a tropical forest in Uganda. Biodiversity and Conservation 15:107–121.

Montgomery RA, Rice KE, Stefanski A, et al. (2020). Phenological responses of temperate and boreal trees to warming depend on ambient spring temperatures, leaf habit, and geographic range. Proceedings of the National Academy of Sciences USA 117:10397–10405.

Naik D, Bhat SSG, Ghate SD, Mustak MS, Rao RSP (2021). Ecological interactions: Patterns of host utilization by tropical butterflies. bioRxiv 474530 (doi:10.1101/2021.12.30.474530).

Naik D, Rao RSP, Kunte K, Mustak MS (2022). Ecological monitoring and indicator taxa: Butterfly communities in heterogeneous landscapes of the Western Ghats and Malabar coast, India. Journal of Insect Conservation 26:107–119.

Navarro-Cano JA, Karlsson B, Posledovich D, et al. (2015). Climate change, phenology, and butterfly host plant utilization. Ambio 44:78–88.

New TR (1997) Are lepidoptera an effective ‘umbrella group’ for biodiversity conservation? Journal of Insect Conservation 1:5–12.

Nitin R, Balakrishnan VC, Churi PV, Kalesh S, Prakash S, Kunte K (2018). Larval host plants of the butterflies of the Western Ghats, India. Journal of Threatened Taxa 10:11495–11550.

Niyogi D, Kishtawal C, Tripathi S, Govindaraju RS (2010). Observational evidence that agricultural intensification and land use change may be reducing the Indian summer monsoon rainfall. Water Resources Research 46:W03533.

Padhye AD, Dahanukar N, Paingankar M, Deshpande M, Deshpande D (2006). Season and landscape-wise distribution of butterflies in Tamhini, northern Western Ghats, India. Zoos’ Print Journal 21:2175–2181.

Padhye AD, Shelke S, Dahanukar N (2012). Distribution and composition of butterfly species along the latitudinal and habitat gradients of the Western Ghats of India. Check List 8:1196–1215.

Parmesan C, Ryrholm N, Stefanescu C, et al. (1999). Poleward shifts in geographical ranges of butterfly species associated with regional warming. Nature 399:579–583.

Pateman RM, Hill JK, Roy DB et al. (2012). Temperature-dependent alterations in host use drive rapid range expansion in a butterfly. Science 336:1028–1030.

Paul M, Sultana A (2020). Studies on butterfly (Insecta: Lepidoptera) diversity across different urban landscapes of Delhi, India. Current Science 118:819–827.

Pinheiro F, Diniz IR, Coelho D, Bandeira MPS (2002). Seasonal pattern of insect abundance in Brazilian Cerrado. Austral Ecology 27:132–136.

Pollard E (1977). A method for assessing changes in the abundance of butterflies. Biological Conservation 12:115–134.

Pozo C, Luis-Martínez A, Llorente-Bousquets J, Salas-Suárez N, Maya-Martínez A, Vargas-Fernández I, Warren AD (2008). Seasonality and phenology of the butterflies (Lepidoptera: Papilionoidea and Hesperioidea) of Mexico’s Calakmul region. Florida Entomologist 91:407–422.

Prudic KL, McFarland KP, Oliver JC, Hutchinson RA, Long EC, Kerr JT, Larrivée M (2017). eButterfly: Leveraging massive online citizen science for butterfly conservation. Insects 8:53.

Purvis A, Hector A (2000) Getting the measure of biodiversity. Nature 40:212–219.

Rao RSP, Girish MKS (2007). Road kills: Assessing insect casualties using flagship taxon. Current Science 92:830–837.

Rákosy L, Schmitt T (2011). Are butterflies and moths suitable ecological indicator systems for restoration measures of semi-natural calcareous grassland habitats? Ecological Indicators 11:1040–1045.

Revathy VS, Mathew G (2013). Seasonality of Rhopalocera (Lepidoptera) species in the butterfly garden at Nilambur in Kerala, southern India. Colemania 35:1–9.

Ribeiro DB, Freitas AVL (2011). Large-sized insects show stronger seasonality than small-sized ones: A case study of fruit-feeding butterflies. Biological Journal of the Linnean Society 104:820–827.

Roy DB, Rothery P, Moss D, Pollard E, Thomas JA (2001). Butterfly numbers and weather: Predicting historical trends in abundance and the future effects of climate change. Journal of Animal Ecology 70:201–217.

Schmitt T, Ulrich W, Delic A, Teucher M, Habel JC (2021). Seasonality and landscape characteristics impact species community structure and temporal dynamics of East African butterflies. Scientific Report 11:15103.

Schroeder PJ, Jenkins DG (2018). How robust are popular beta diversity indices to sampling error? Ecosphere e02100.

Sengupta P, Banerjee KK, Ghorai N (2014). Seasonal diversity of butterflies and their larval food plants in the surroundings of upper Neora Valley National Park, a sub-tropical broad leaved hill forest in the eastern Himalayan landscape, West Bengal, India. Journal of Threatened Taxa 6:5327–5342.

Shapiro AM (1976). Seasonal polyphenism. In: Hecht MK, Steere WC, Wallace B (eds) Evolutionary Biology. Springer, Boston, MA pp 259–333.

Sharma K, Acharya BK, Sharma G, Valente D, Pasimeni MR, Petrosillo I, Selvan T (2020). Land use effect on butterfly alpha and beta diversity in the Eastern Himalaya, India. Ecological Indicators 110:105605.

Sharma N, Sharma S (2021). Assemblages and seasonal patterns in butterflies across different ecosystems in a sub-tropical zone of Jammu Shiwaliks, Jammu and Kashmir, India. Tropical Ecology 62:261–278.

Shimadzu H, Dornelas M, Henderson PA, Magurran AE (2013). Diversity is maintained by seasonal variation in species abundance. BMC Biology 11:98.

Singh AP, Gogoi L, Sebastain J (2015). The seasonality of butterflies in a semi-evergreen forest: Gibbon Wildlife Sanctuary, Assam, northeastern India. Journal of Threatened Taxa 7:6774–6787.

Sokal RR, Rohlf FJ (1995). Biometry (3rd ed.). WH Freeman, New York. 887 pages.

Sreekumar PG, Balakrishnan M (2001). Habitat and altitude preferences of butterflies in Aralam Wildlife Sanctuary, Kerala. Tropical Ecology 42:277–281.

Suman A, Ravikanthachari N, Kunte K (2021). A comparison between time-constrained counts and line transects as methods to estimate butterfly diversity in tropical habitats. bioRxiv https://doi.org/10.1101/2021.09.04.458959.

Tiple A, Agashe D, Khurad AM, Kunte K (2009). Population dynamics and seasonal polyphenism of Chilades pandava butterfly (Lycanidae) in Central India. Current Science 97:1774–1779.

Tiple AD, Khurad AM (2009). Butterfly species diversity, habitats and seasonal distribution in and around Nagpur city, central India. World Journal of Zoology 4:153–162.

van Bergen E, Osbaldeston D, Kodandaramaiah U, et al. (2017). Conserved patterns of integrated developmental plasticity in a group of polyphenic tropical butterflies. BMC Evolutionary Biology 17:59.

Varikoden H, Revadekar JV, Kuttippurath J, Babu CA (2019). Contrasting trends in southwest monsoon rainfall over the Western Ghats region of India. Climate Dynamics 52:4557–4566.

Warren MS, Hill JK, Thomas JA, et al. (2001). Rapid responses of British butterflies to opposing forces of climate and habitat change. Nature 414:65–69.

