## Supplemental for "Seasonal dynamics and polyphenism of butterfly communities in the coastal plains of central Western Ghats, India"

Running title: Seasonality of butterflies of the Western Ghats.

*Keywords: Biodiversity loss, Climate change, Eco-informatics, Ecological monitoring, Habitat degradation, Polyphenism and seasonality*

### Supplemental Materials and Methods

#### *Study sites*

In the study region (the Western Ghats), four months from June to September receive heavy rain (~85% of annual total, Fig. S1) and are considered as monsoon period (IMD, https://mausam.imd.gov.in/). Other months can be categorized into post-monsoon (October to January) and pre-monsoon (February to May) periods. The monsoon and post-monsoon periods can be considered as wet seasons, whereas pre-monsoon period which receives least amount of rain (less than 6%) as dry season. Where appropriate, seasonality of butterflies was explored based on these distinct periods/seasons.

### Supplemental Results and Discussion

#### *Inter-annual variation*

There were noticeable inter-annual variations both in rain fall (Fig. S1) and butterfly species compositions (Fig. 1). For example, rainfall in 2018 was 226% and 33% more in pre-monsoon and monsoon periods, respectively, compared to the previous year. On the contrary, butterfly abundance in 2018 was 34% and 24% less in pre-monsoon and monsoon-periods, respectively, compared to the previous year! For example, the overall abundance of the most abundant butterfly *Euploea* spp. was 29.1% less in 2018 compared to the previous year and it was significantly different from the overall decrease (22.2%) in the abundance of butterflies in the same period ($p = 0.026$, Z-test for two-proportions). The abundance of the second most abundant butterfly *Catopsilia* spp. was striking - nearly 75.8% less compared to the previous year. However, given the short-term nature of this study, inter-annual differences were not considered further.

#### *Seasonal distribution of singletons and low-abundant species*

28 species of butterflies were sighted only in one season (six in pre-monsoon, 12 in monsoon, and 10 in post-monsoon). These were mostly very low-abundant species. For example, there were 18 singleton species, and except for *Graphium antiphates* (n = 127), the number of individuals in other species were four or less. Another set of 28 low-abundant species were sighted only in two seasons, while the remaining 119 species were present in all three seasons (Table S1).

#### *Seasonal polyphenism*

At least 66 species (out of 336 species) in the Western Ghats are known to display some form of seasonal pholyphenism (Bhakare and Ogale, 2018; Kunte *et al.*, 2021; Nitin *et al.*, 2018). As many as 48 out of 175 species identified in this study from the Western Ghats are known to have seasonal polyphenism (Table S1; Bhakare and Ogale, 2018). However, only five species which could easily/clearly be identified as DSF or WSF were explored in detail here. The remaining polyphenic species were not explored for various reasons such as (a) difficulty of identification in the field, (b) lack of clear difference between DSF and WSF, (c) low abundance, and (d) absence of polyphenism in the given condition/habitat. For example, while *Ypthima huebneri* (N = 2249) and *Eurema hecabe* (n = 2143) were similarly abundant, the latter was very difficult to differentiate in to DSF and WSF in the field (Fig. S6A and B). On the other hand, *Junonia lemonias* was both less abundant (n = 205), and DSF and WSF (based on underside wing patterns) were difficult to distinguish in the field as the species mostly sits with open wings (Fig. S6C).

**Table S1.** Monthly/seasonal patterns of butterflies of the Western Ghats (Table S1.xlsx).

**Table S2.** Distribution pattern of number of species and individuals of butterflies in different (taxonomic) families and seasons.

| | **Species** | | | | | **Individuals** | | | | | **Individuals[#]** | | | | **Individuals[$]** | | |
|---|---|---|---|---|---|---|---|---|---|---|---|---|---|---|---|---|---|
| | **Pre-monsoon** | **Monsoon** | **Post-monsoon** | **Total** | **$\chi^2$ (p-value)[+]** | **Pre-monsoon** | **Monsoon** | **Post-monsoon** | **Total** | **$\chi^2$ (p-value)[+]** | **Pre-monsoon** | **Monsoon** | **Post-monsoon** | **Total** | **Pre-monsoon** | **Monsoon** | **Post-monsoon** |
| **Hesperiidae** | 22 | 34 | 28 | 39 | 0.19 (0.66) | 142 | 1032 | 1084 | 2258 | 110.2 (9E-26) | 0.01 | 0.09 | 0.05 | 0.05 | 0.06 | 0.46 | 0.48 |
| **Lycaenidae** | 40 | 37 | 45 | 49 | 0.0056 (0.94) | 1727 | 1339 | 4896 | 7962 | 359.3 (4E-80) | 0.16 | 0.12 | 0.23 | 0.18 | 0.22 | 0.17 | 0.61 |
| **Nymphalidae** | 49 | 48 | 50 | 54 | 0.29 (0.59) | 5616 | 5612 | 11200 | 22428 | 6.0 (0.014) | 0.52 | 0.50 | 0.53 | 0.52 | 0.25 | 0.25 | 0.50 |
| **Papilionidae** | 16 | 16 | 16 | 17 | 0.11 (0.74) | 883 | 1364 | 1127 | 3374 | 150.3 (2E-34) | 0.08 | 0.12 | 0.05 | 0.08 | 0.26 | 0.40 | 0.33 |
| **Pieridae** | 11 | 13 | 14 | 15 | 0.11 (0.74) | 2403 | 1889 | 2781 | 7073 | 430.1 (2E-95) | 0.22 | 0.17 | 0.13 | 0.16 | 0.34 | 0.27 | 0.39 |
| **Riodinidae** | 0 | 1 | 1 | 1 | 0.65 (0.42) | 0 | 8 | 15 | 23 | 5.7 (0.017) | 0 | 0 | 0 | 0 | 0 | 0.35 | 0.65 |
| **Total** | 138 | 149 | 154 | 175 | | 10771 | 11244 | 21103 | 43118 | | | | | | 0.25 | 0.26 | 0.49 |

[+]The chi-squared test for trend in proportions for species/individuals against total in three (pre-monsoon, monsoon, and post-monsoon) seasons. Seasonal trends in species richness within families were not significantly different from the total. Being the most abundant in terms of individuals, Nymphalidae had a seasonal trend more similar to the total, but other families had very dissimilar seasonal trends.
[#]Family-wise and [$]season-wise proportions.

**Table S3.** Monthly/seasonal variations in the diversity indices[#] of butterfly communities of the Western Ghats.

| | **Pre-monsoon** | | | | | **Monsoon** | | | | | **Post-monsoon** | | | | | **Total** |
|---|---|---|---|---|---|---|---|---|---|---|---|---|---|---|---|---|
| | **Feb** | **Mar** | **Apr** | **May** | **Total** | **Jun** | **Jul** | **Aug** | **Sep** | **Total** | **Oct** | **Nov** | **Dec** | **Jan** | **Total** | |
| Number of individuals | 2701 | 2137 | 1878 | 4055 | 10771 | 2168 | 2080 | 2471 | 4525 | 11244 | 5816 | 6308 | 5054 | 3925 | 21103 | 43118 |
| Number of species (richness/α-diversity) | 109 | 96 | 94 | 105 | 138 | 106 | 107 | 104 | 134 | 149 | 135 | 129 | 132 | 121 | 154 | 175 |
| Rarefied richness (α-diversity) | 100.1 | 93.7 | 93.8 | 88.5 | 137.7 | 102.9 | 104.5 | 99.1 | 117.8 | 148.3 | 113.0 | 110.0 | 109.1 | 104.7 | 146.6 | 153.7 |
| Shannon's H' (α-diversity) | 3.65 | 3.67 | 3.61 | 2.87 | 3.61 | 3.83 | 3.77 | 3.91 | 4.07 | 4.03 | 4.03 | 3.97 | 3.85 | 3.76 | 4.04 | 4.05 |
| Simpson's 1-D (α-diversity) | 0.95 | 0.96 | 0.94 | 0.82 | 0.94 | 0.97 | 0.96 | 0.97 | 0.97 | 0.97 | 0.97 | 0.97 | 0.97 | 0.96 | 0.97 | 0.97 |
| Fisher's (α-diversity) | 22.8 | 20.7 | 20.8 | 19.7 | 22.3 | 23.3 | 23.9 | 22.0 | 25.9 | 24.3 | 24.7 | 23.0 | 24.8 | 23.6 | 22.5 | 23.3 |
| Brillouin (α-diversity) | 3.57 | 3.59 | 3.52 | 2.82 | 3.58 | 3.74 | 3.68 | 3.83 | 4.00 | 3.99 | 3.98 | 3.93 | 3.80 | 3.70 | 4.02 | 4.04 |
| Whittaker's $\beta_W$ (β-diversity)[+] | 0.21 | 0.23 | 0.25 | 0.22 | 0.13^ | 0.22 | 0.22 | 0.24 | 0.20 | 0.13^ | 0.18 | 0.19 | 0.18 | 0.18 | 0.12^ | 0.21 |
| Dominance D (dominance) | 0.05 | 0.04 | 0.06 | 0.18 | 0.06 | 0.03 | 0.04 | 0.03 | 0.03 | 0.03 | 0.03 | 0.03 | 0.03 | 0.04 | 0.03 | 0.03 |
| Berger-Parker (dominance) | 0.14 | 0.14 | 0.20 | 0.39 | 0.16 | 0.09 | 0.08 | 0.08 | 0.09 | 0.08 | 0.08 | 0.08 | 0.10 | 0.10 | 0.07 | 0.07 |
| Menhinick (richness) | 2.10 | 2.08 | 2.17 | 1.65 | 1.33 | 2.28 | 2.35 | 2.09 | 1.99 | 1.41 | 1.77 | 1.62 | 1.86 | 1.93 | 1.06 | 0.84 |
| Margalef (richness) | 13.7 | 12.4 | 12.3 | 12.5 | 14.8 | 13.7 | 13.9 | 13.2 | 15.8 | 15.9 | 15.5 | 14.6 | 15.4 | 14.5 | 15.4 | 16.3 |
| Chao-1 (richness) | 132.0 | 105.2 | 111.0 | 124.1 | 157.0 | 121.8 | 124.8 | 112.0 | 138.6 | 155.1 | 144.2 | 134.0 | 157.3 | 138.5 | 158.5 | 196.9 |
| Pielou's J' (evenness/equitability) | 0.78 | 0.80 | 0.79 | 0.62 | 0.73 | 0.82 | 0.81 | 0.84 | 0.83 | 0.81 | 0.82 | 0.82 | 0.79 | 0.78 | 0.80 | 0.78 |
| Buzas-Gibson (evenness/equitability) | 0.35 | 0.41 | 0.39 | 0.17 | 0.27 | 0.43 | 0.41 | 0.48 | 0.44 | 0.38 | 0.42 | 0.41 | 0.36 | 0.35 | 0.37 | 0.33 |

[#]As obtained using Past (version 3.26) software (https://past.en.lo4d.com/windows) (see Naik *et al*., 2022).
[+]Mean β-diversity against all other months (^or seasons).

**Table S4.** Indicator values for butterfly species based on month/seasonality (Table S4.xlsx).

**Table S5.** Seasonality of polyphenic species.

| | | Pre-monsoon | Monsoon | Post-monsoon | Total | $\chi^2$ (p-value)[+] |
|---|---|---|---|---|---|---|
| *Jamides celeno* | WSF | 53 | 195 | 926 | 1174 | 371.2 (1E-82) |
| | DSF | 179 | 0 | 122 | 301 | |
| | Total | 232 | 195 | 1048 | 1475 | |
| *Melanitis leda* | WSF | 2 | 28 | 53 | 83 | 7.7 (0.0055) |
| | DSF | 24 | 2 | 42 | 68 | |
| | Total | 26 | 30 | 95 | 151 | |
| *Mycalesis* spp.[#] | WSF | 5 | 176 | 140 | 321 | 0.35 (0.55) |
| | DSF | 33 | 0 | 73 | 106 | |
| | Total | 38 | 176 | 213 | 427 | |
| *Ypthima baldus* | WSF | 82 | 402 | 645 | 1129 | 138.3 (6E-32) |
| | DSF | 96 | 0 | 61 | 157 | |
| | Total | 178 | 402 | 706 | 1286 | |
| *Ypthima huebneri* | WSF | 209 | 344 | 1423 | 1976 | 495.2 (1E-109) |
| | DSF | 207 | 0 | 66 | 273 | |
| | Total | 416 | 344 | 1489 | 2249 | |
| Total | WSF | 351 | 1145 | 3187 | 4683 | 852.4 (2E-187) |
| | DSF | 539 | 2 | 364 | 905 | |
| | Total | 890 | 1147 | 3551 | 5588 | |

[#]Mostly *M. mineus*.

[+]The chi-squared test for trend in proportions for WSF versus DSF in three (pre-monsoon, monsoon, and post-monsoon) seasons. In *Melanitis leda and more strongly in Mycalesis* spp., there was a shift in the WSF toward monsoon and DSW toward post-monsoon as compared to other species wherein WSF was predominant in post-monsoon and DSF in pre-monsoon.

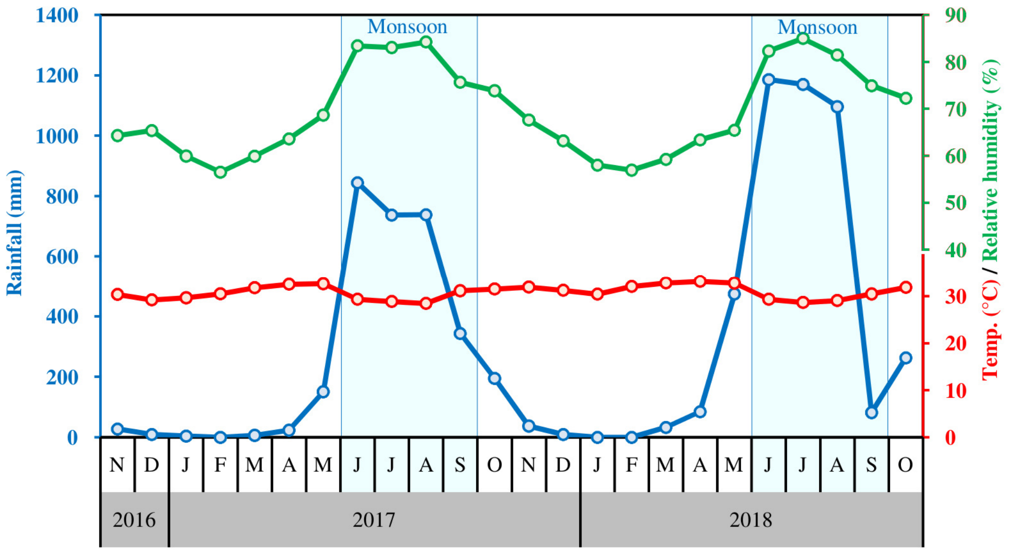


**Fig. S1.** Rainfall (mm), mean day temperature (°C), and relative humidity (%) in the study region (Dakshina Kannada district) during the study period between Nov 2016 and Oct 2018. It rains heavily during June to Sept (monsoon season); however, there is a clear difference in the amount of rain between 2017 and 2018. Overall, the region shows high humidity (>57%), which further increases noticeably during the monsoon. The mean day temperature is around 30°C which shows only a slight dip during the monsoon.

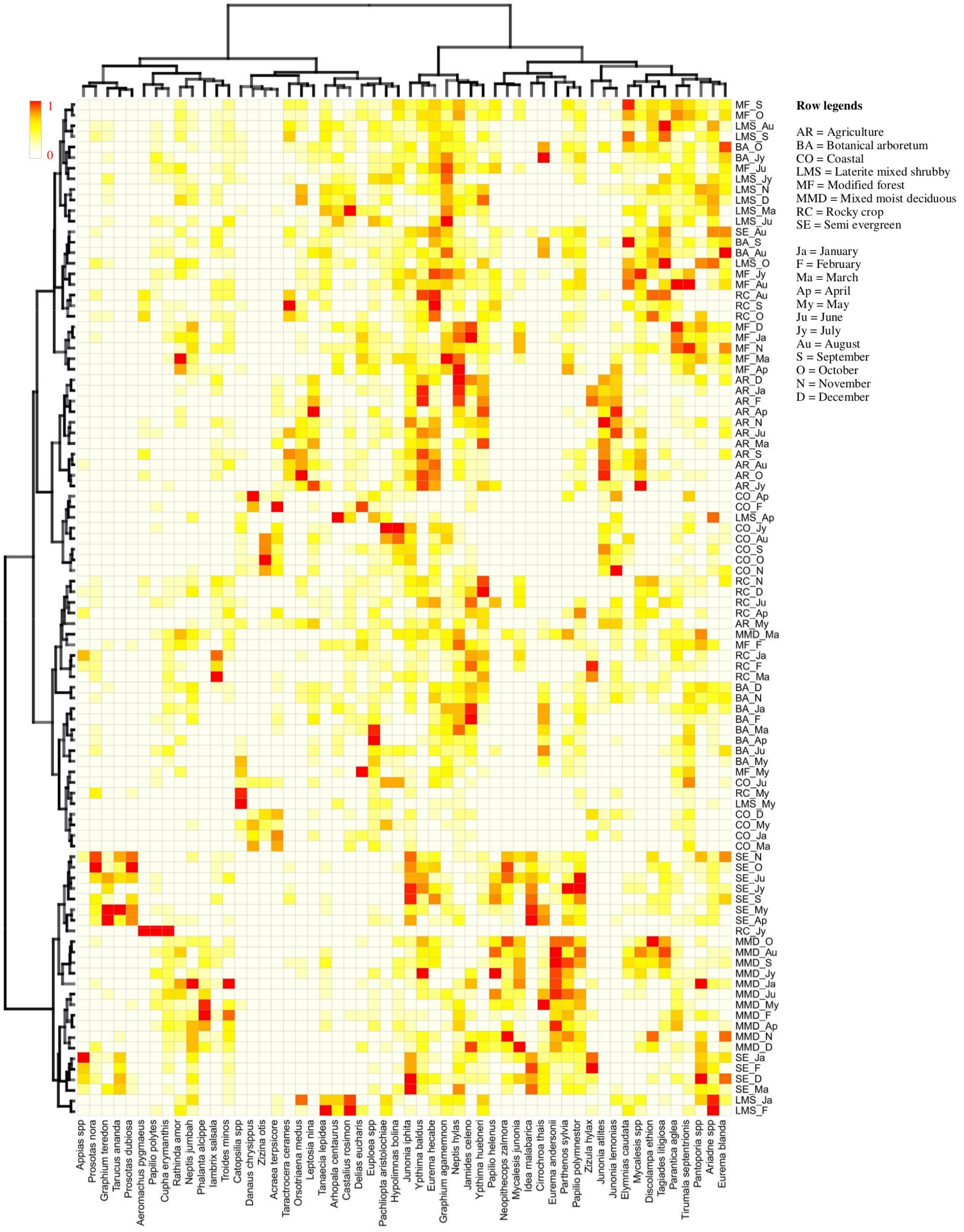


**Fig. S2.** Heat map and HCA of seasonality (months) versus site-specificity of butterflies (54 species that together make up more than 90% of the total abundance) in the coastal plains of the central Western Ghats. Samples mostly cluster based on sites (habitats) as different habitats show large differences in the abundance of species. For example, most of the samples from agriculture (AR) or mixed moist deciduous (MMD) habitats cluster together.

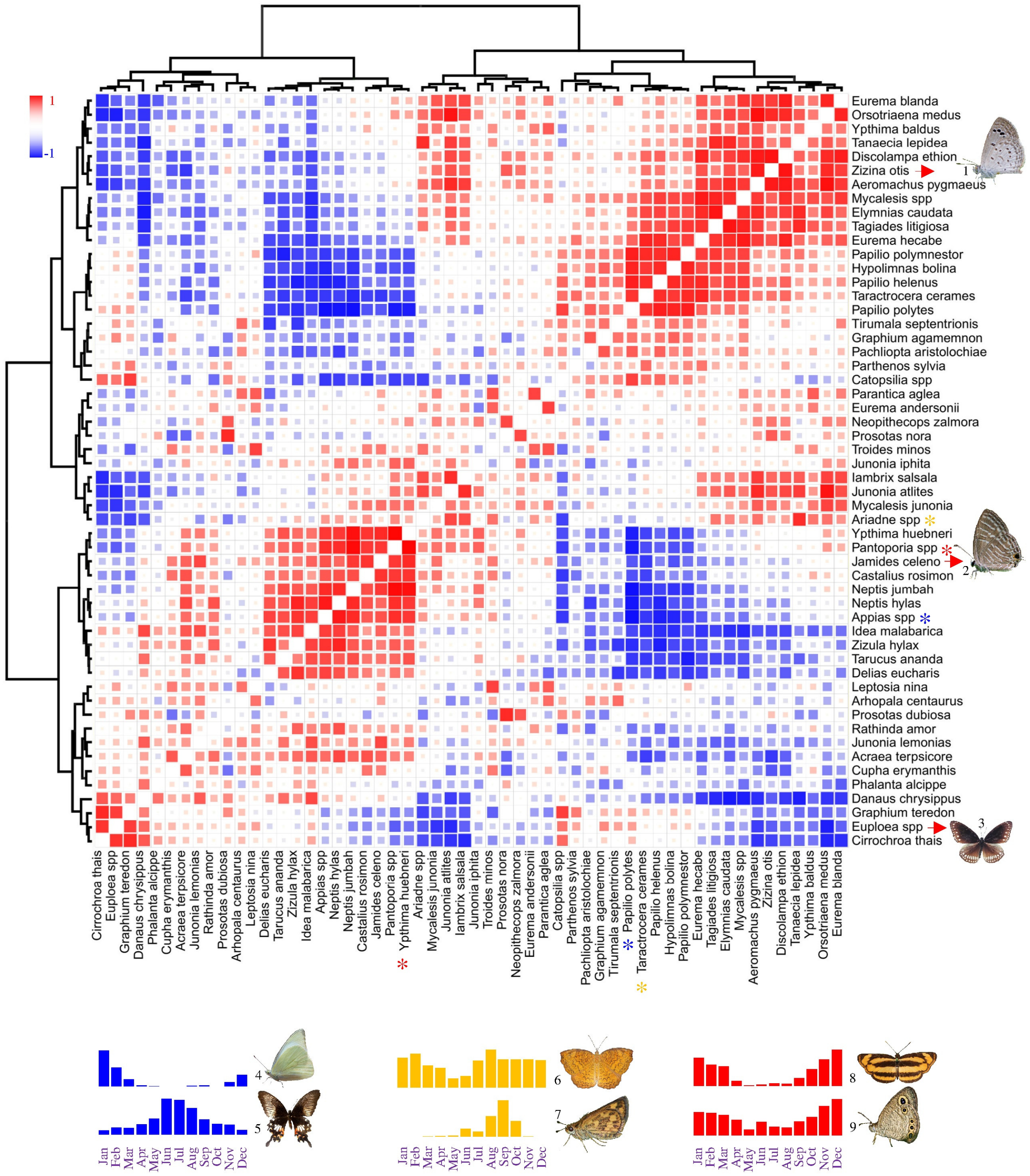


**Fig. S3.** Spearman's rank correlation plot of seasonality of butterflies (54 species that together make up more than 90% of the total abundance) in the coastal plains of the central Western Ghats. Several species-clusters can be seen based on the distinct seasonality patterns. For example, cluster which includes *Zizina otis* [1], peaks in October, *Jamides celeno* [2] in December-January, and *Euploea* spp. [3] in April. Non-diagonal correlation ranges from -0.928 ($p = 1.3E-5$, *Appias* spp. [4] versus *Papilio polytes* [5]) to 0.003 ($p \approx 1$, *Ariadne* spp. [6] versus *Taractrocera ceramas* [7]) to 0.965 ($p = 3.8E-7$, *Pantoporia* spp. [8] versus *Ypthima huebneri* [9]).

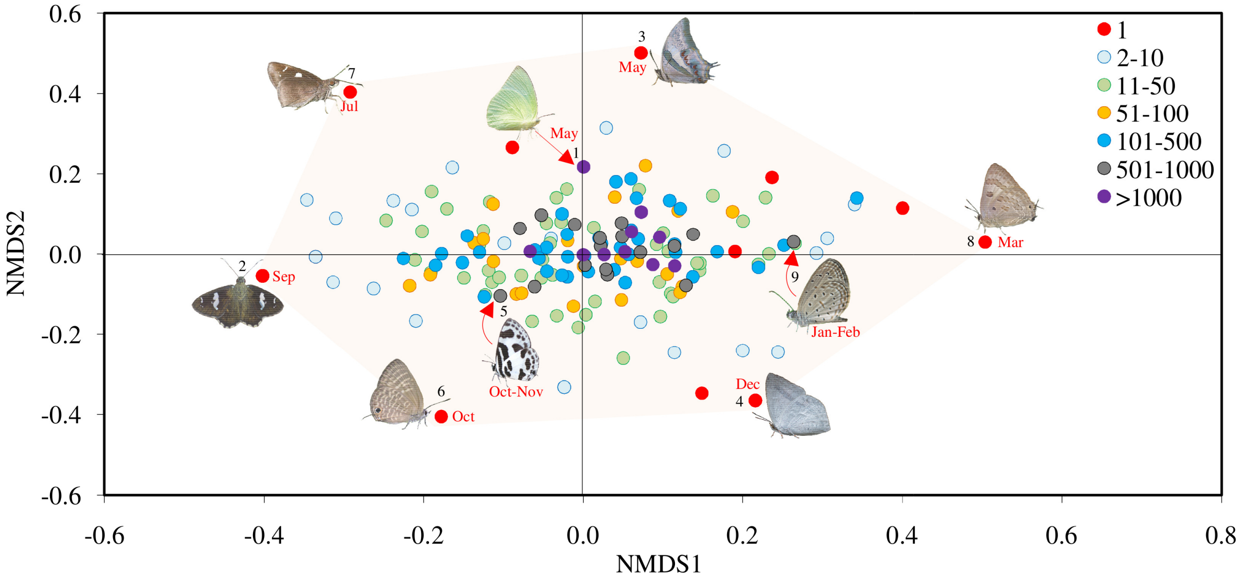


**Fig. S4.** Species clustering using NMDS. The NMDS plot (stress = 0.12) showed a pattern based on the abundance of species. The species that were very abundant clustered closely together at the center. Less abundant species are scattered, and the singletons are at the periphery. Flagged species in the figure are (1) *Catopsilia* spp., (2) *Celaenorrhinus putra*, (3) *Charaxes schreiber*, (4) *Curetis siva*, (5) *Discolampa ethion*, (6) *Nacaduba pactolus*, (7) *Notocrypta curvifascia*, (8) *Virachola perse*, and (9) *Zizula hylax*.

| | Jan | Feb | Mar | Apr | May | Jun | Jul | Aug | Sep | Oct | Nov | Dec |
|---|---|---|---|---|---|---|---|---|---|---|---|---|
| | | Pre-monsoon | | | | Monsoon | | | | Post-monsoon | | |
| *Graphium antiphates* | 0 | 0.34 | 0.78 | 0.41 | 0.19 | 0 | 0 | 0 | 0 | 0 | 0 | 0 |
| *Kallima horsfieldii* | 0 | 0 | 0 | 0 | 0.07 | 0.13 | 0.37 | 0.52 | 0.52 | 0.16 | 0 | 0 |
| *Doleschallia bisaltide* | 0 | 0 | 0 | 0 | 0.12 | 0.12 | 0.25 | 0 | 0.55 | 0 | 0.12 | 0.12 |
| *Belenois aurota* | 0 | 0 | 0 | 0 | 0 | 0.13 | 0 | 0.13 | 0.13 | 0.68 | 0 | 0 |
| *Zeltus amasa* | 0 | 0 | 0 | 0 | 0 | 0.11 | 0 | 0 | 0 | 0.16 | 0.67 | 0.28 |
| *Anthene lycaenina* | 0 | 0.11 | 0 | 0.15 | 0.09 | 0 | 0 | 0 | 0 | 0.41 | 0.66 | 0.16 |
| *Badamia exclamationis* | 0.15 | 0.07 | 0.18 | 0 | 0 | 0.10 | 0 | 0.07 | 0.15 | 0.15 | 0.46 | 0.46 |
| *Anthene emolus* | 0.32 | 0 | 0 | 0 | 0 | 0.13 | 0 | 0 | 0.09 | 0 | 0.48 | 0.39 |
| *Rohana parisatis* | 0 | 0.09 | 0 | 0 | 0 | 0 | 0 | 0.09 | 0.19 | 0.32 | 0.50 | 0.32 |
| *Notocrypta paralysos* | 0.10 | 0 | 0 | 0 | 0 | 0 | 0 | 0.14 | 0.14 | 0.33 | 0.24 | 0.33 |
| *Lethe europa* | 0.20 | 0.10 | 0 | 0 | 0 | 0.10 | 0 | 0 | 0.20 | 0.49 | 0.17 | 0.14 |
| *Graphium doson* | 0 | 0 | 0 | 0.14 | 0.45 | 0.14 | 0.10 | 0.14 | 0.43 | 0.17 | 0 | 0.07 |
| *Appias spp* | 0.68 | 0.37 | 0.23 | 0.06 | 0.06 | 0 | 0 | 0.03 | 0.10 | 0.02 | 0.30 | 0.44 |
| *Lampides boeticus* | 0.49 | 0.44 | 0.44 | 0.25 | 0 | 0 | 0 | 0 | 0.03 | 0.19 | 0.15 | 0.44 |
| *Zizula hylax* | 0.63 | 0.63 | 0.24 | 0.09 | 0.01 | 0 | 0 | 0 | 0.03 | 0.04 | 0 | 0.34 |
| *Spindasis lohita* | 0.31 | 0.11 | 0.27 | 0.11 | 0.11 | 0 | 0 | 0 | 0 | 0 | 0.11 | 0.33 |
| *Catochrysops strabo* | 0.30 | 0.23 | 0.08 | 0.10 | 0.42 | 0.18 | 0 | 0 | 0 | 0.08 | 0 | 0.31 |
| *Eurema laeta* | 0.37 | 0.35 | 0.29 | 0.14 | 0.35 | 0 | 0.12 | 0 | 0 | 0 | 0.16 | 0.19 |
| *Rapala manea* | 0.46 | 0.15 | 0.08 | 0.15 | 0 | 0.08 | 0 | 0 | 0.15 | 0.11 | 0.35 | 0.42 |

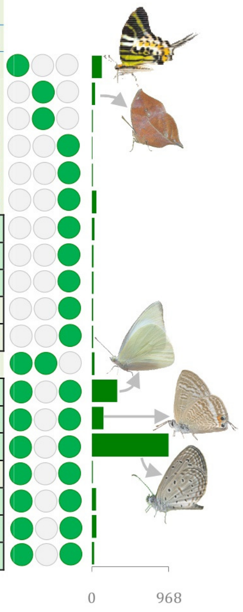


**Fig S5.** Indicator species. Just 11 single-season-specific indicator species were found based on indicator value estimation (numbers in green are significant [when taken on monthly or seasonal basis] at $p < 0.05$, permutation test; De Cáceres *et al*., 2010). While there were numerous species common to monsoon and post-monsoon seasons (see Table S4), only a very few indicator species were present in combination with pre-monsoon (dry) season. Filled green circles indicate the seasonal (pre-monsoon, monsoon, and post-monsoon) presence. Bar diagram at the right shows the absolute abundance of species. See Table S4 for indicator values of species at different combinations of months or seasons.

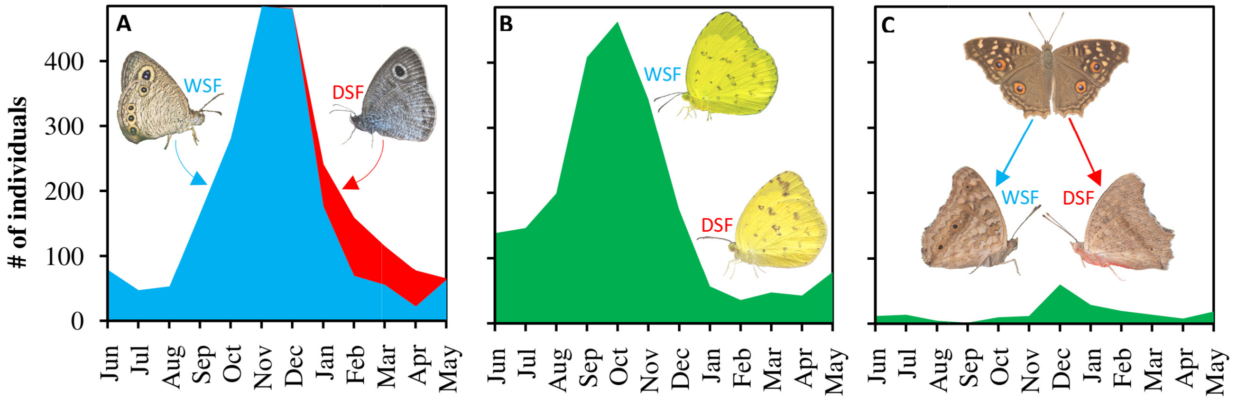


**Fig. S6.** Polyphenism of butterfly species in the Western Ghats. Seasonal distributions of (A) WSF and DSF in *Ypthima huebneri* (N = 2249). (B) *Eurema hecabe* (n = 2143) - WSF and DSF are difficult to differentiate in the field. (C) *Junonia lemonias* (n = 205) - sample size is very less, and WSF and DSF (based on underside wing patterns) are difficult to distinguish in the field as the species mostly sits with open wings. See Table S1 for polyphenic species identified in this study in the Western Ghats. Note that the wing colour difference between WSF and DSF in *Ypthima huebneri* was possibly due to camera/lighting issue in the field.
